# In-silico Screening of *Origanum vulgare* Phytocompounds as Potential Drug Agents Against *Vp35* Protein of the Ebola Virus

**DOI:** 10.1101/2023.03.25.534218

**Authors:** Malaika Saeed, Aqeela Ashraf, Burhan Sabir, Muhammad Osama Zafar, Muhammad Hassan Raza, Rashid Saif

## Abstract

The leading cause of the Ebola virus outbreak during 2013-16 in Western Africa was a lack of targeted anti- viral drug choices, a fast rate of mutations and the unavailability of many of the structural proteins and annotations within its genome. The surroundings of the Ebola River in DR-Congo fail to get rid of this endemic, the reason behind this was believed to be its origin from non-human primates, which made its risk assessment and tracing difficult. The Vp35 is a multifunctional protein with innate immune antagonistic properties and is considered one of the most suitable drug targets within this virus. The main motive of this study is to discover a potential anti-viral drug against the Ebola virus by targeting the aforementioned protein with different phytocompounds of oregano that have the lowest binding energies and qualifies over different simulation parameters, so firstly, molecular docking was performed on its 28 compounds using PyRx to get the best complexes with minimum binding energies e.g., -8.9Kcal/mol. Ligands with the best docking scores were gone through Lipinski’s rule of five for drug likeliness potential. For the drug affirmation, molecular dynamic simulation was also performed with the best two docked complexes using NAMD/VMD to find out their conformational stability through RMSD, RMSF, Rg, SASA and H-bond analyses. Current computer-generated prediction suggested that Ursolic acid and Oleanolic acid possess potential inhibitory effects against virus replication. Furthermore, paradigm shifts of usage of natural and herbal products for treating infectious diseases are being encouraged here, However, further wet-lab experiments and clinical trials are still needed to determine the robustness of these virtually tested phytocompounds against the Vp35 protein of the Ebola virus.

## 1. Introduction

Ebola Virus Disease (EVD) or Ebola Hemorrhagic Fever (EHF) is caused by the Ebola virus, affecting humans and various primates [1]. This virus is a negative-sense RNA virus, and classified under the filoviridae family [2,3]. As the infection progresses, it damages the immune system and multiple organs, resulting in a drop in blood-coagulating cells and which leads to swear bleeding. The outbreak of 1976 caused the emergence of EHF, claiming the lives of up to 90% of infected individuals [4, 5].

Africa has seen the most cases and occurrences of the Ebola Virus Disease [5]. The 2014-2016 Ebola outbreak in West Africa initially emerged in southeastern Guinea and quickly spread to metropolitan areas and across borders [6,7]. The exact origin of the Ebola virus is unknown, but based on similar viruses, it is believed to be animal-borne with bats or nonhuman primates being the most likely sources of transmission for EVD [8]. Infected animals carrying the virus can transmit it to other animals, such as chimps, monkeys, gorillas, and ultimately, to the human population [9].

The Ebola virus is known to specifically target liver cells, immune system cells, and endothelial cells of blood vessels by producing glycoproteins that facilitate macropinocytosis [10]. By damaging the endothelial cells, the virus causes increased permeability of blood vessels and loss of cellular adhesion. Additionally, infection of liver cells can lead to compromised detoxification abilities and further damage to the immune system. These mechanisms contribute to the rapid spread of the Ebola virus and the extensive damage [4]. The Ebola virus is transmitted to humans through direct contact with infected animals and can be spread through bodily fluids of infected individuals or contaminated objects. The virus enters the body through damaged skin or mucous membranes and can also be transmitted through sexual contact, as it can persist in certain bodily fluids after recovery [11].

Vp35 is a protein in the Ebola virus that plays a critical role in the virus’s replication process. It functions as a suppressor of the immune system’s response to the virus, allowing the virus to evade detection and destruction by the host’s immune system. Vp35 also helps to regulate viral gene expression, ensuring that the virus is able to replicate efficiently. Because of its importance in the virus’s life cycle, Vp35 has been identified as a potential target for antiviral therapies aimed at inhibiting Ebola virus replication.

Furthermore, the lack of a specific treatment for Ebola virus, exploring antiviral properties of herbal compounds may offer a potential solution. *Origanum vulgare*, for instance, contains phytochemicals that have been demonstrated to exhibit antiviral activity and have shown effectiveness against various viruses in experimental studies.

## 2. Materials and Methods

### 2.1 Selection of medicinal herb (Origanum vulgare)

*Origanum vulgare* is an important plant and has anti-viral abilities. It has many compounds (Figure 1) that show antiviral properties while quercetin in it is the most important and can bind the glycoproteins on the viral envelope [12]. It has 60 compounds present in it, Carvacrol and Thymol are the major ones in it. They are cultivated in European countries and are used for flavoring [13]. The chemical properties of the 28 ligands were described in Table 1.

**Table 1.**
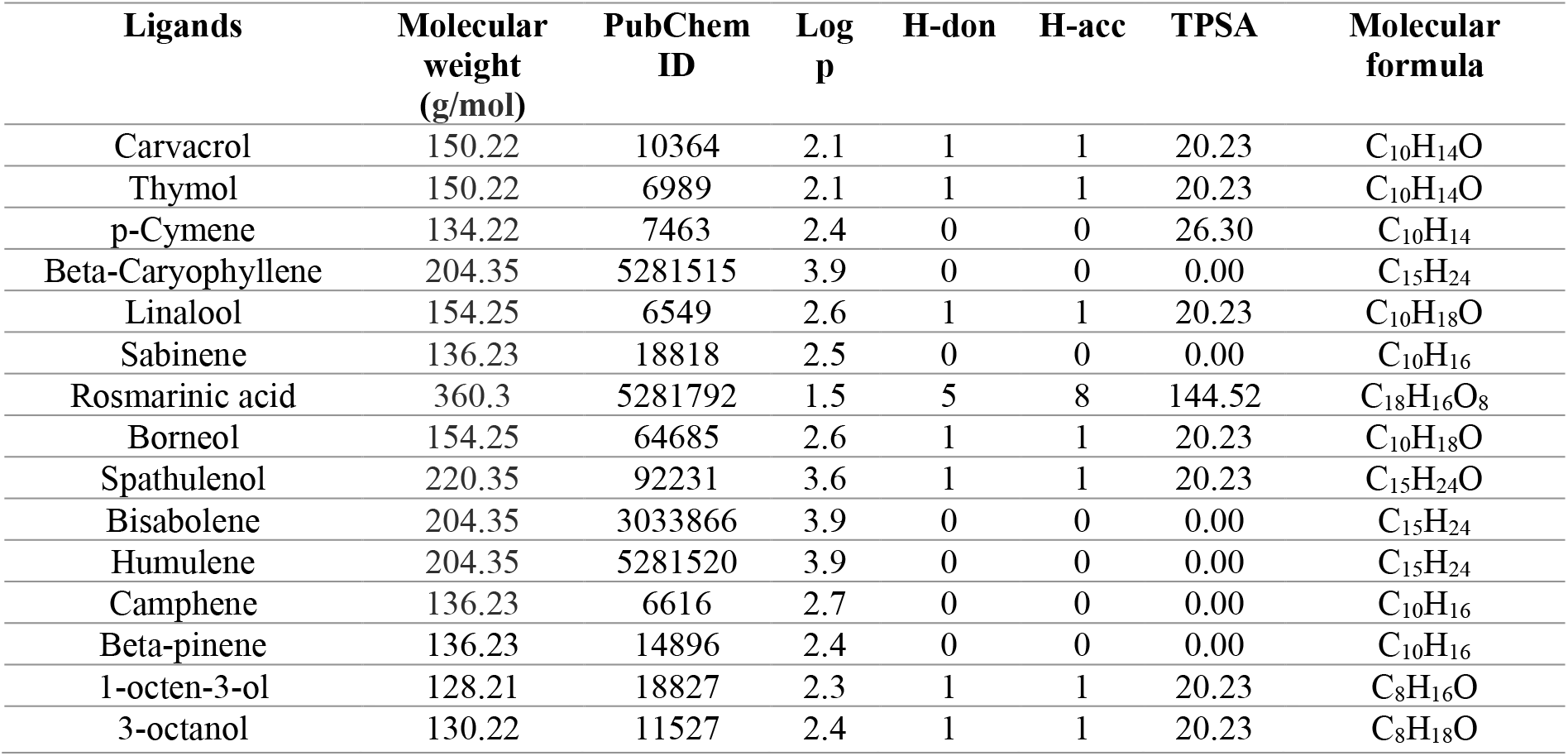
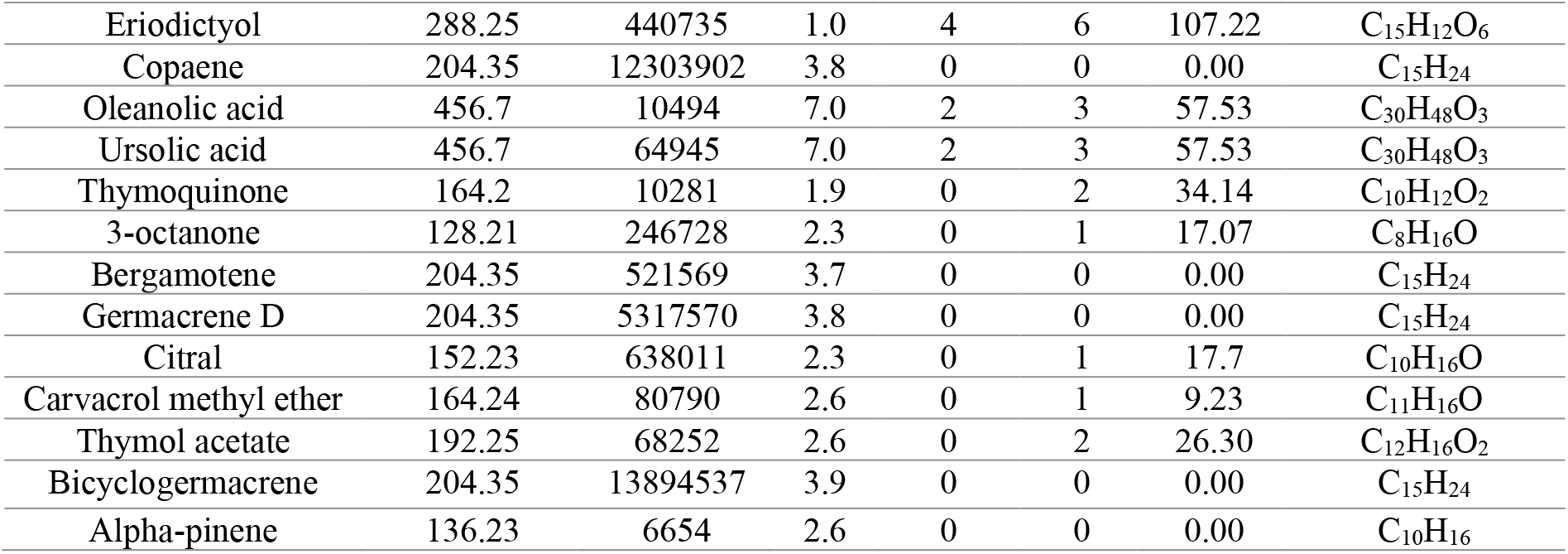
Chemical and structural properties of different phytocompounds of *Origanum vulgare*

**Figure 1.**
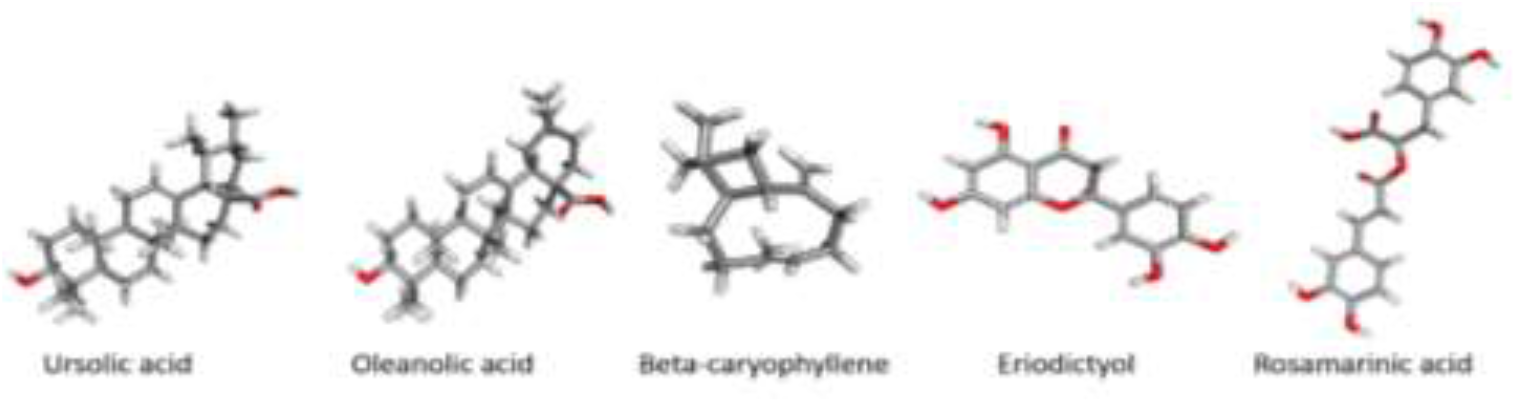
Chemical structures of few of the ligands of *Origanum vulgare*

**Figure 2.**
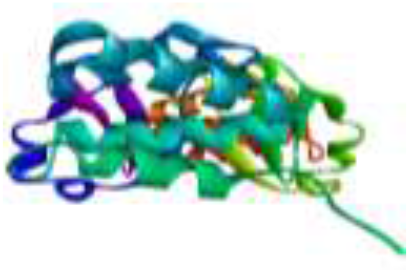
Prepared Vp35 (PDB ID:3FKE) Ebola virus protein

### 2.2 Selection of targeted protein

Vp35, a fundamental viral cofactor of RNA-dependent RNA polymerase, that is crucial for Ebola virus replication and host intrinsic immune emission. Ebola virus replication is regulated by the phosphorylation of Vp35. Thus, by targeting Vp35, the replication process can be halted [14]. The crystallographic properties of Vp35 protein as described in Table 2.

**Table 2.** Crystallographic properties of Vp35

| Protein | PDB ID | Classification | Virus | Expression system | Method | Resolution | Atom count | Structural weight | Chains |
| --- | --- | --- | --- | --- | --- | --- | --- | --- | --- |
| Vp35 | 3FKE | RNA binding protein | Ebola virus | E. Coli | X-ray diffraction | 1.4Å | 2143 | 28.42kda | A, B |

### 2.3 Molecular docking by PyRx

PyRx was used for molecular docking, and it is a drug discovery software that enables the analysis of protein-ligand interactions for drug-like properties. It provides a comprehensive package that includes result visualization, modeling, design simulations, and process optimization.

#### 2.3.1 Preparation of ligand

Numerous databases such as ChEMBL, Zinc, Merck, PubChem, Enamine, DrugBank, Asinex, etc. are accessible to get the required ligands in SDF format. These selected ligands were prepared using PyRx by uploading the sdf format file and make it a macromolecule by removing water molecules and already attached ligands.

#### 2.3.2 Preparation of protein

A Protein Data Bank (PDB) was used to download targeted protein with the PDB ID of 3FKE. It was prepared using discovery studio software (DS) by applying following steps i.e., removal of water molecules, removal of inhibitors and repeated chains to make the active site available for a new ligand. Similarly, water molecules were eliminated from the protein surface with the goal that the active site wouldn’t be concealed during docking process.

### 2.4 Docking process

Ligand-protein docking aims to predict the optimal binding configuration of a ligand with a known three-dimensional structure of a protein. To achieve this, the protein and ligand molecules were prepared and uploaded in the pdbqt format onto the main interface of PyRx software. Energy minimization was performed by selecting the AMBER force field for 500 steps to create the protein-ligand complex. Subsequently, the Autodock Vina wizard command was utilized to execute the docking process.

### 2.5 Pfizer rule of five for drug likeliness

The Lipinski’s (Pfizer) rule was applied on the phytocompounds/ligands of *Origanum vulgare* by AdmetSAR and Swiss ADME to determine the pharmacological properties.

### 2.6 Molecular dynamic simulation using VMD/NAMD softwares

The molecular dynamics simulation was performed to investigate the microscopic interaction dynamics, structural deviation, fluctuation, and conformational changes and the thermodynamics of protein-ligand complex. VMD and NAMD softwares were used simultaneously for efficient scaled simulation of a bimolecular large system, running on a computer processors or graphic processing unit (GPU) cores. The stability of the best docked complex of Oleanolic Acid and Ursolic Acid with Vp35 was analyzed by plotting histograms of RMSD, RMSF, SASA, Rg, H-bonds using simulation results.

#### 2.6.1 Complex parameterization & simulation process

The hydrogen atoms missing in the ligand structure were added using Open Babel and the resulting structure was converted into a mol2 file format. This file was then parameterized using the CHARMM-GUI input generator. Similarly, the protein structure files were generated using VMD autopsf plugins. A script was written for complex formation and executed on the command line interface. A solvation box was created to provide a medium for interaction between the complex system. Molecular dynamics simulations were run for 1 nanosecond (5000000 steps) and energy minimization was done using the conjugate gradient method. The periodic boundary conditions were established for the energy-minimized complex to run the simulation. A constant temperature of 310K and pressure of 1 atm were maintained during the simulation process. After adjusting all the parameters, the MD simulation was executed using NAMD/VMD software.

## 3. Results

### 3.1 Docking scores of Origanum vulgare compounds

The docking of Vp35 was performed with 28 compounds of *Origanum vulgare* to find out the best binding energies. The docking scores of all the phytocompounds were reported in table 3 in which the Oleanolic acid and Ursolic acid have the highest binding energies.

**Table 3.** Docking scores of *Origanum vulgare* with 3FKE protein

| Ligands | Docking score |
| --- | --- |
| Carvacrol | -5.4 |
| Thymol | -5.3 |
| p-Cymene | -5.5 |
| Caryophyllene | -6.4 |
| Linalool | -4.7 |
| Sabinene | -4.7 |
| Rosmarinic acid | -7.0 |
| Borneol | -5.7 |
| Spathulenol | -6.6 |
| Bisabolene | -5.9 |
| Humulene | -6.4 |
| Camphene | -4.8 |
| Beta-pinene | -4.9 |
| 1-octen-3-ol | -3.8 |
| 3-octanol | -3.9 |
| Eriodictyol | -7.1 |
| Copaene | -6.4 |
| Oleanolic Acid | -8.9 |
| Ursolic Acid | -8.9 |
| Thymoquinone | -5.1 |
| 3-octanone | -3.7 |
| Bergamotene | -5.7 |
| Germacrene D | -6.5 |
| Citral | -4.5 |
| Carvacrol methyl ether | -5.9 |
| Thymol acetate | -5.9 |
| Bicyclogermacrene | -6.5 |
| Alpha-pinene | -4.9 |

### 3.2 2D/3D interactions of best docked complexes

Moreover, the 2D/3D interactions of the best docked complex which have the binding affinities of -8.9 were depicted in Figure 3. In Ursolic acid-Vp35 complex, one conventional hydrogen bond with amino acid: glutamine (GLN A:329) is formed, 10 alkali bonds with amino acids: PRO A:316, PRO A:318, VAL A:327, VAL A:314, VAL A:294, ALA B:291, ALA B:290, LEU B:249 are present, 6 Van der Waals interaction of amino acids: ILE B:286, PRO B:293, THR A:335, HIS A:296, VAL B:294, PRO B:292 are in the surroundings. Whereas, In Oleanolic acid a single conventional hydrogen bond with amino acid: proline (PRO B:292) is formed, and 10 alkali bonds with amino acids: PRO B:292, PRO B:318, VAL B:327, VAL A:314, VAL B:314, VAL B:294, ALA B:291, ALA A:291, LEU A:249 are present, 12 Van der Waals interaction of amino acids: VAL A:294, PRO B:316, PRO B:315, PRO B:319, ALA A:290, ILE A:286, SER A:253, ASP A:252, THR B:335, GLN B:329, PRO A:292, SER B:317 are in the surroundings.

**Figure 3.**
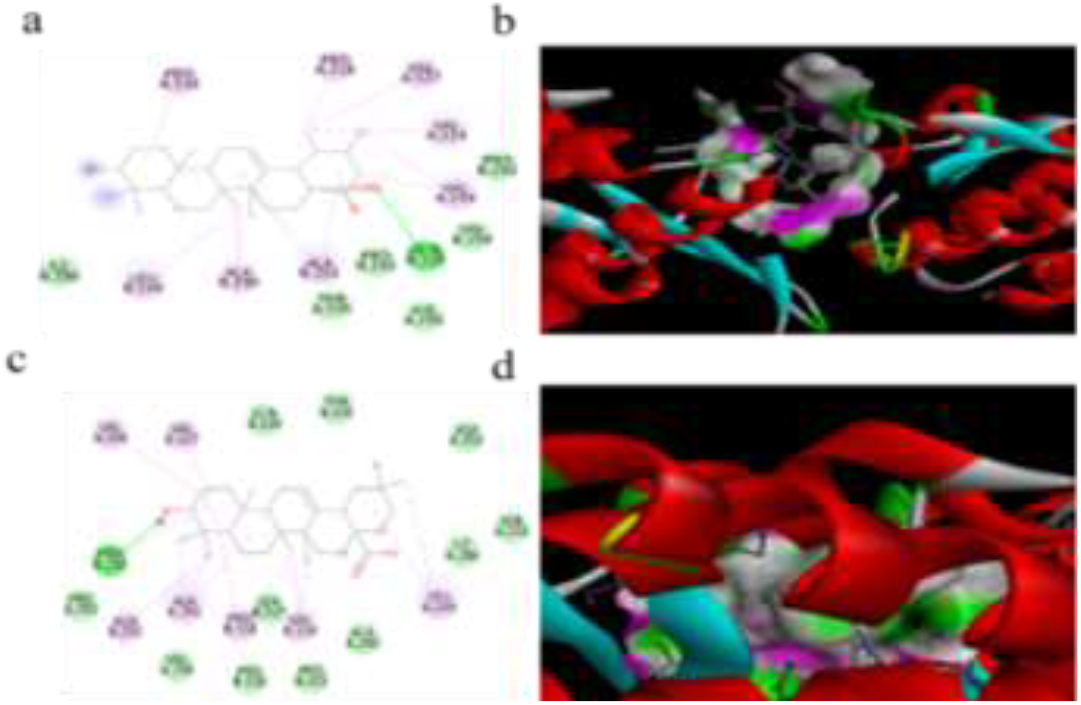
2D/3D interactions of Ursolic Acid (a & b), 2D**/**3D interactions of Oleanolic Acid (c & d)

### 3.3 Lipinski’s rule outcomes of ligands

Lipinski’s rule of 5 states that a ligand satisfying a minimum of two out of the five defined criteria is a favorable candidate for drug development [16]. To assess the pharmacological properties of the top docked ligands, the SwissADME tool was employed, adhering to Lipinski’s guidelines as delineated in Table 4.

**Table 4.** Drug likeliness attributes of top five oregano phytocompounds

| PubChem ID | Ligands | Mol. weight (<500 Da) | LogP (<5) | H-donor | H-acceptor | Violations | Docking scores |
| --- | --- | --- | --- | --- | --- | --- | --- |
| 10494 | Oleanolic acid | 456 | 7.0 | 2 | 3 | 1 | -8.9 |
| 64945 | Ursolic acid | 456 | 7.0 | 2 | 3 | 1 | -8.9 |
| 440735 | Eriodictyol | 288 | 1.0 | 4 | 6 | 0 | -7.4 |
| 5281792 | Rosmarinic acid | 360 | 1.5 | 5 | 8 | 0 | -7.0 |
| 5317570 | Germacrene D | 204 | 3.8 | 0 | 0 | 1 | -6.5 |

### 3.4 Post simulation analysis for ligands-3FKE protein complex

#### 3.4.1 Root mean square deviation (RMSD) analysis

Computations were performed to measure the typical distance between the group of molecules. After completion of the simulation process of ligand-protein complex. The RMSD values were determined for every setup in a given direction for every design of a similar direction utilizing the standard RMSD trajectory tool of VMD. The RMSD values ranging between 2-3Å are normal, if it exceeds 4Å then it may indicate significant problems with the protein folding process [17]. The RMSD values of Ursolic acid and Oleanolic acid were ranges from 2.5Å to 4Å as shown in the Figure 4 which indicated that the RMSD value of both the complexes were deviated and shifted towards slightly unstable state.

**Figure 4.**
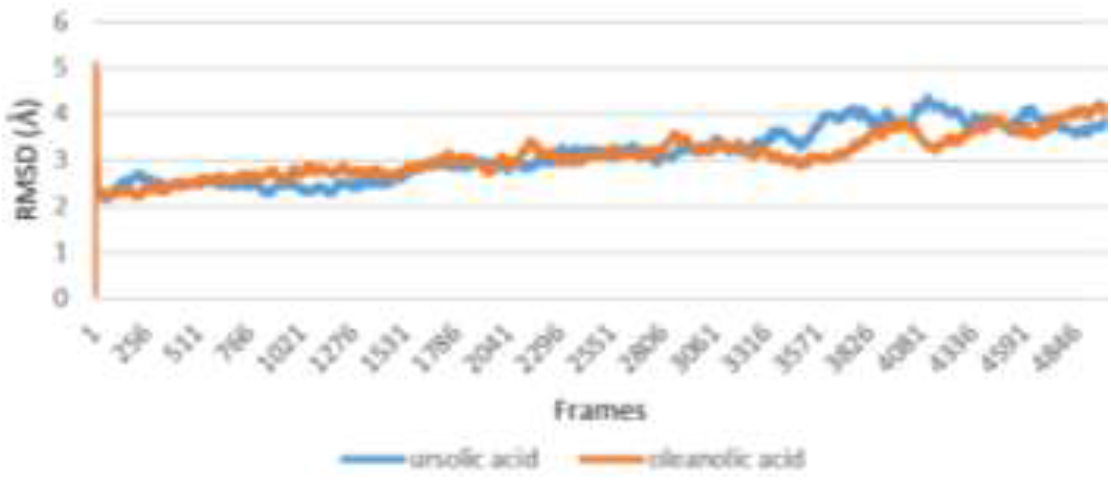
RMSD plots of Ursolic Acid & Oleanolic Acid complexes with Vp35

#### 3.4.2 Root mean square fluctuations (RMSF) analysis

The estimation of RMSF was performed to portray the proportion of vacillation of ligand-protein structures in the binding since the limiting postures, energy, and interaction changeability relies upon a residual variance. The RMSF is the time average of the RMSD, a mathematical estimation only like of RMSD however the individualized computation of residue adaptability is estimated as opposed to zeroing in on contrasts between the entire structure over time. The ideal value of RMSF depends on 3Å very much like RMSD, 3.4Å is acceptable too [18]. Both ligands show abnormality at different points in Figure 5. Initially Oleanolic acid was stable for approximately 125 frames and then its value raised to 7.9Å and after that, it went to normal values ranging from 1- 2.3Å. In the case of Ursolic acid initially, it was crossing the normal range of RMSF values but from frame 55 to 195 it was fluctuating from 1.5-4 Å and then onwards it got normal.

**Figure 5.**
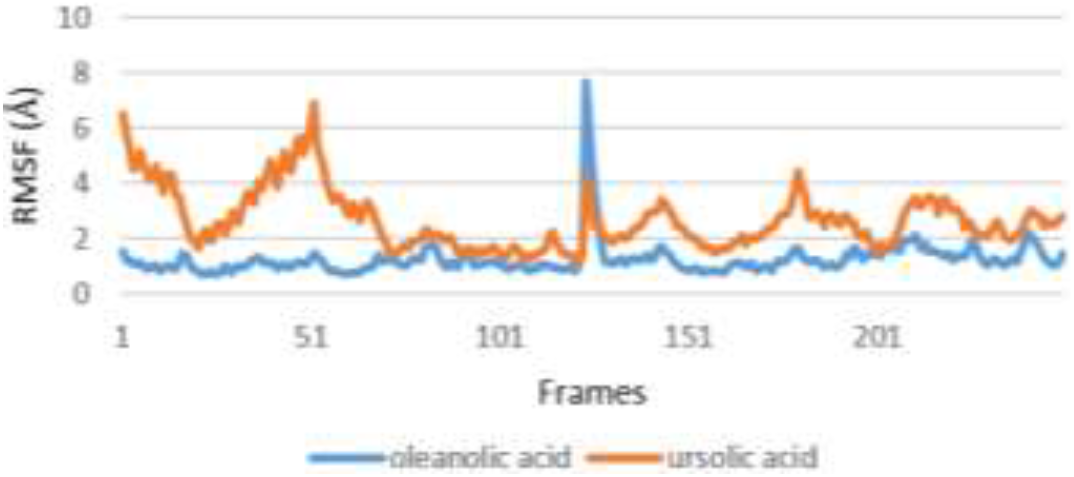
RMSF plots of Ursolic Acid & Oleanolic Acid complexes with Vp35

#### 3.4.3 Hydrogen bonding

The hydrogen bond analysis was carried out to determine the number of hydrogen bonds and their manageability in the framework. Complex buildups will generally be flawless as the gap between the residues is decreased with the formation of hydrogen bonding between the molecules. Hydrogen bonds in the dynamic simulation are distinguished by; donor-acceptor distance (3Å or less), and donor- hydrogen-acceptor point (150° or more). Both ligands have shown the compact graphs (Figure 6) in the following descriptive manner; major compounds formed by Ursolic acid that stabilized the Ursolic–Vp35 complex makes 116.7% occupancy rate with ARG298 acceptor and PHE328 donor with an occupancy rate of 63.49% respectively. Similarly, SER299 in Oleanolic acid was proved to stabilize the Oleanolic-Vp35 complex with the occupancy rate of 69.45%.

**Figure 6.**
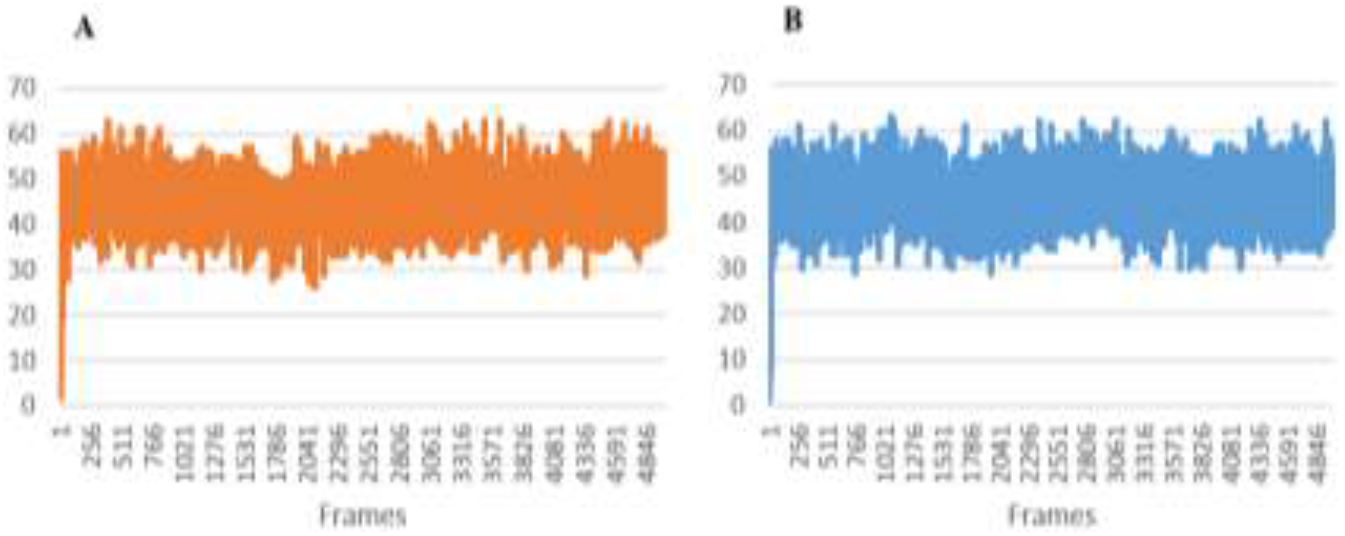
Hydrogen bond analysis **A)** Oleanolic Acid **B)** Ursolic Acid

#### 3.4.4 Solvent accessible surface area (SASA) analysis

The SASA analysis is based on protein surface area characterized around a hypothetical centre of solvent particle where van der waals interacts with its molecule. It is considered to be a conjunctive factor in declaring protein structure, folding and stability. Backbone atoms of the complex can be illustrated by assessing the SASA parameters, structure compactness is indicated if the value of protein solvent accessible surface area is low. A SASA value of Oleanolic and Ursolic acids were in 15000 nm^2^ range as shown in Figure 7, which suggested that a significant portion of the protein’s surface is exposed and accessible to solvent molecules, indicating a more flexible or dynamic protein structure.

**Figure 7.**
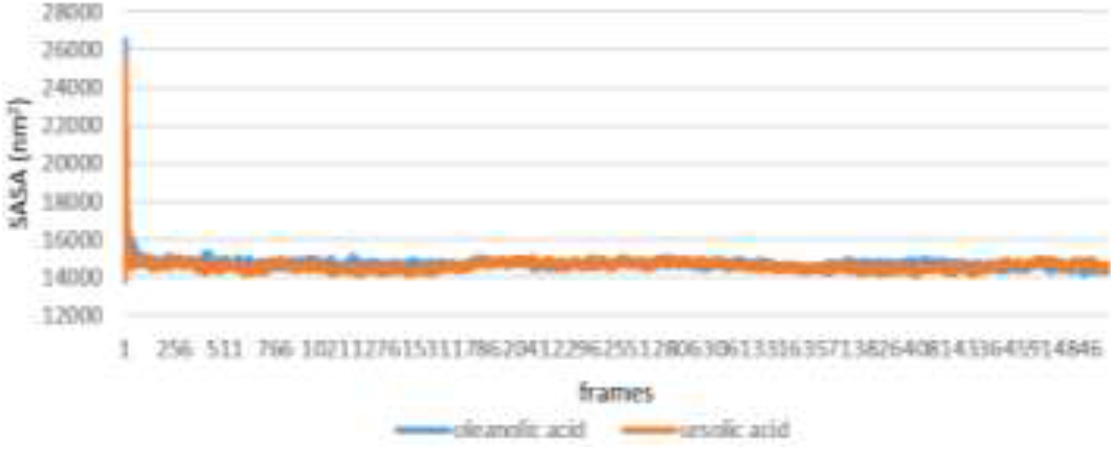
SASA plots of Ursolic Acid & Oleanolic Acid complexes with Vp35

#### 3.4.5 Radius of gyration (Rg) analysis

The radius of gyration is the distribution of amino acids around its axis, the analysis fundamentally indicates chain molecule size, and is calculated to measure the variations of ligand/protein complex during MD simulations. The Rg value evaluates the flexibility and compactness manner of protein while it is attached to the phytocompounds, so the hydrodynamic radius of the protein structure per time can be monitored. The Rg values of the Ursolic and Oleanolic Acid complexes were ranging from 21.6Å to 20.8Å which indicated that the complexes were slightly differ from the overall stability range of Rg values (Figure 8).

**Figure 8.**
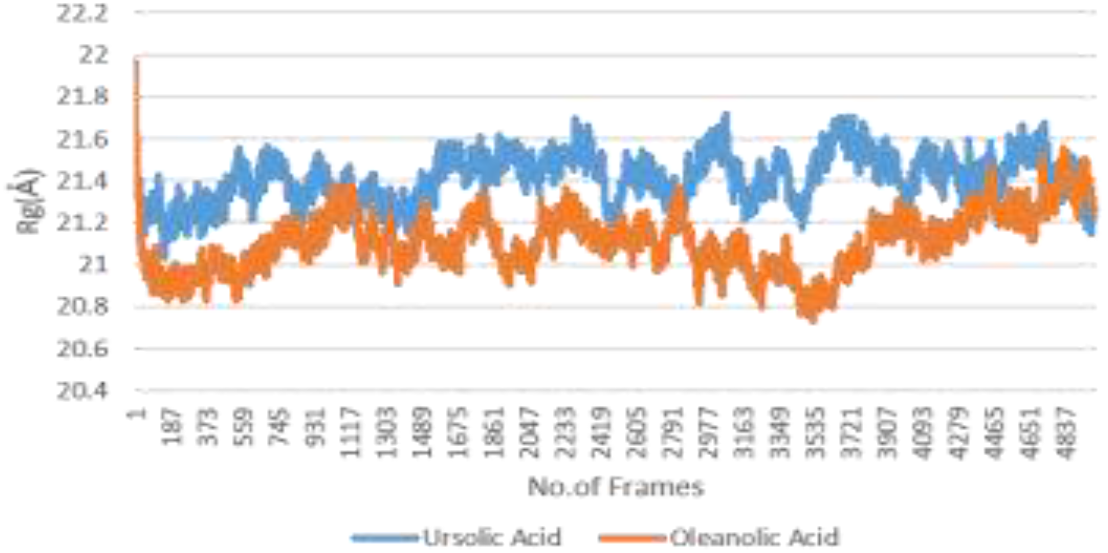
Rg plots of Ursolic Acid & Oleanolic Acid complexes with Vp35

## 4. Discussion

Commercial drugs that demonstrated efficacy in combating the Ebola virus also causes adverse side effects. First of all, clinical trials for these drugs have not been conducted e.g., Fevipiravir. Secondly, there are risk factors i.e., these drugs can induce teratogenicity and embryo toxicity [19]. During the 2014-2016 Ebola epidemic in West Africa, a few clinical trials were conducted to assess the safety and efficacy of remdesivir in treating Ebola virus disease. These trials found that remdesivir showed some promise in treating Ebola, however further research is needed to establish its efficacy.

Oleanolic acid and Ursolic acid, derived from *Origanum vulgare*, demonstrated the lowest binding energy during docking and molecular dynamic simulations, suggesting stable formation of the protein-ligand complex by optimally fitting into the active site of 3FKE and engaging in a maximum number of hydrogen bonds. These results are noteworthy, as these compounds exhibit potential as inhibitors of the Ebola virus, based on the observed protein-ligand interactions and stability attained through computational drug design via molecular docking and molecular dynamic simulation.

Inmazeb is currently the available drug for treating EVD, with its compounds showing docking scores ranging from -8.0 to -8.8Kcal/mol according to the literature [20]. However, this medication is associated with side effects such as chills, faster heartbeat, vomiting, and tachypnea [21]. Oleanolic acid and Ursolic acid were both subjected to post-simulation analysis, and their RMSD values suggest that they have the potential to be suitable drugs against the Ebola virus protein Vp35. Oleanolic acid showed better results than Ursolic acid, with more stable RMSF values. Swiss ADME analysis confirmed that both compounds are non-toxic and can be used without any harmful effects. Ursolic acid is already in use for the treatment of Rotavirus. It is a pentacyclic triterpenoid with antiviral abilities that has been extensively studied in vitro and in vivo [22]. Similarly, Oleanolic acid is also an antiviral compound that is considered a novel drug in the treatment of HIV and influenza, as well as other viruses [23]. Both of these drugs are also used in the treatment of Hepatitis C, as they inhibit NS5B activity [24].

Due to the high pathogenicity of the Ebola virus and its classification as one of the deadliest viruses by the WHO, there is a critical need to find a specific drug against EVD. Oleanolic acid and Ursolic acid can be considered potential inhibitors against the Ebola virus as they follow Lipinski’s rules of drug likeliness. Moreover, if simulations are run for a longer time, they have the potential to demonstrate more stability.

## Conclusion

This study aimed to identify potential Ebola inhibitors from Oregano by targeting the Vp35 protein essential for viral replication. Molecular docking and dynamic simulation revealed Oleanolic and Ursolic acids derived from *Origanum vulgare* to have the lowest binding energies and exhibit stable complex formation, suggesting their potential as effective Ebola inhibitors. Further investigation through research and clinical trials is crucial to assess the effectiveness of these replication inhibitors in managing Ebola virus outbreaks in future.

